# CLEAN: Leveraging spatial autocorrelation in neuroimaging data in clusterwise inference

**DOI:** 10.1101/2022.03.02.482664

**Authors:** Jun Young Park, Mark Fiecas

**Author notes:** Corresponding author, *Email address:* (Jun Young Park).

## Abstract

While clusterwise inference is a popular approach in neuroimaging that improves sensitivity, current methods do not account for explicit spatial autocorrelations because most use univariate test statistics to construct cluster-extent statistics. Failure to account for such dependencies could result in decreased reproducibility. To address methodological and computational challenges, we propose a new powerful and fast statistical method called CLEAN (**C**lusterwise inference **Le**veraging spatial **A**utocorrelations in **N**euroimaging). CLEAN computes multivariate test statistics by modelling brain-wise spatial autocorrelations, constructs cluster-extent test statistics, and applies a refitting-free resampling approach to control false positives. We validate CLEAN using simulations and applications to the Human Connectome Project. This novel method provides a new direction in neuroimaging that paces with advances in high-resolution MRI data which contains a substantial amount of spatial autocorrelation.

## 1. Introduction

The Human Connectome Project (HCP) has incorporated state-of-the-art advances in neuroimaging acquisition and processing [1]. In particular, the analytic approach using surface-based analysis has attractive features compared to volumetric analysis, including dimension reduction and improved cross-subject alignment and visualization [2, 3, 4]. Surface-based data has been used in `FreeSurfer` [5] and is available as a `CIFTI` format developed by the HCP for the analysis of the grayordinate data [6]. The `CIFTI` format is available for processing, analysis, or visualization in `Python` [2], `R` [7], and `Matlab` [8].

Statistical methods for surface-based analysis, however, are still lacking, and there remain a number of challenges that need to be addressed. First, surface-based data exhibit strong spatial autocorrelation that impacts the sensitivity and specificity of statistical tests. Second, cluster-wise inference with adjustments for multiple comparisons can lead to the use of a stringent threshold, further affecting sensitivity and specificity [9]. Finally, the use of small sample size (e.g. *N* ≤ 30) has been shown to impact the reproducibility of the results [10, 11]. Current statistical practices include spatial smoothing (blurring) of the images to reduce the noise variance at each location which could improve statistical power. However, spatial smoothing decreases the benefits gained from having high-resolution data, but it has also been shown to potentially lead to a significant loss in sensitivity and specificity [3].

We first review existing statistical methods that improve statistical power in neuroimaging data with spatial autocorrelations in Sections 1.1 and 1.2 [12]. It is summarized into two major areas: including modelling the mean (i.e. regression parameters in the general linear model (GLM)) and modelling the covariance (i.e. noise in GLM). We then provide an outline of our approach in 1.3.

### 1.1. Modelling the mean structure: clusterwise inference

In this paper, we define clusterwise inference as a statistical method that aggregates test statistics (e.g. *z* or *t* statistics) in the spatial domain [13]. Clusterwise inference has good statistical power when signal locations form a spatial cluster within the brain [14, 15]. Two approaches include (i) a two-step approach, where first an initial threshold for each location to define spatial clusters is set and then inference on the clusters is conducted and (ii) a one-step approach that constructs a cluster-enhanced statistic for every location and set a threshold that controls the familywise error rate (FWER). The former approach relies on setting an initial cluster-defining threshold (CDT) that balances false positives with power. While *p* < 0.001 is commonly used [16], setting a model-based or adaptive threshold has also been proposed at the expense of computational cost [17, 18]. The latter approach is represented by the threshold-free cluster enhancement (TFCE), where the FWER-controlling threshold is determined by either resampling methods or Gaussian Random Field Theory (GRFT) [14, 19]. Other related methods have shown to be powerful in task-fMRI by borrowing information from the neighboring locations through a non-linear filter [15].

Despite its popularity, current clusterwise inference is limited mainly to volumetric (3D) fMRI data or relies heavily on GRFT. Also, a comprehensive exploratory data analysis revealed that the spatial covariance structure is far from Gaussian but closer to exponential or a mixture of Gaussian and exponential, implying that the assumptions needed by GRFT may be too strong, motivating a need for new methods to replace GRFT [20, 21].

While spatial smoothing increases signal-to-noise ratio (SNR) in each location, unwanted variations arise when conducting statistical inference. We consider each location to be either a signal location or a null location. After smoothing, a null location may become a signal location if other signal locations are nearby, and is thus a pseudo-signal since it was truly null prior to smoothing. Thus, any powerful statistical inference that identifies the pseudo-signal location as signal would be prone to decreased specificity. Also, smoothing is not beneficial when a signal cluster (i.e. a collection of neighboring and spatially connected signal locations) is small in size, in which case these small signal clusters may only be detected if they have large effect sizes that would decrease when blurred with neighboring null locations, implying that smoothing can also decrease sensitivity.

### 1.2. Modelling the covariance structure: spatial autocorrelation

Most clusterwise inference methods use univariate test statistics (e.g. voxel-wise *t* statistics) to compute cluster-enhanced test statistics. However, neuroimaging data, especially surface-based data, reveals a high degree of spatial autocorrelation. Ignoring such dependencies in constructing test statistics in each location could result in low sensitivity when the signal-to-noise ratio (SNR) is low. In clusterwise inference, this would be problematic when the size of a signal region is small.

In the neuroimaging literature, there are both parametric and nonparametric approaches to model the spatial structure of the test statistics. Examples of a nonparametric approach are the methods that locally smooth the variance of the test statistics. FSL provides a variance smoothing option [10] to generate ‘pseudo’ *t* statistics, and Wang et al. [22] recently proposed an approach based on empirical Bayes, termed ‘locally moderated’ *t* statistic, by pooling variance information across a predefined set of neighbors. An example of a parametric approach includes modelling the spatial covariance using a Gaussian process (GP). Indeed, this parametric approach is the most straightforward and commonly-used way to address the spatial autocorrelation and reduce the variance of a test statistic at each location. One can find this approach in constructing multivariate test statistics [23, 12, 24] or in constructing a posterior probability map [4]. From the modelling perspective, several autoregressive (AR) models have been proposed in the neuroimaging literature, including the simultaneously autoregressive (SAR) model [25] or the conditional autoregressive (CAR) model [26]. However, while both SAR and CAR models account for spatial autocorrelations, the implied spatial covariance structure by CAR or SAR is often misleading in approximating the spatial covariance [27]. Also, in task-fMRI data in 3D, the parametric structure for the spatial covariance could be prone to misspecification, e.g. Eklund et al. [20] illustrated that the covariance structure is Gaussian locally but exponential in large distances. Another possibility is to construct an explicit spatial covariance using a pairwise distance matrix [23, 12, 24]. However, without any simplification, the spatial Gaussian process is computationally infeasible when the number of spatial locations is large. Therefore, it is common to divide the brain into several homogeneous regions based on atlases and model each region separately [12, 24] or use a data-driven approach [23]. However, these approaches depend heavily on ROI selections or parcellation algorithms and do not benefit from modelling the autocorrelation structure in the entire brain.

We comment that, because fitting a model that accounts for the spatial structure of the data is already computationally intensive, resampling would not be applicable in general to control FWER. Moreover, to our knowledge, its extension to clusterwise inference (modelling mean structure) has not been developed in the neuroimaging literature.

### 1.3. Our contribution: new clusterwise inference leveraging spatial autocorrelation modelling

This paper aims to develop a fast and powerful one-step clusterwise inference method that measures the association between an image and a covariate while explicitly modelling the spatial autocorrelation in the data. We address improving the sensitivity of statistical tests by modelling both mean structures (clusterwise inference) and covariance structures (Gaussian process) simultaneously. While (i) the clusterwise inference with univariate statistics and (ii) massive-univariate inference with multivariate statistics are developed to improve the power of the brain imaging studies, to our best knowledge, there is no method for clusterwise inference with multivariate test statistics that adjusts for spatial autocorrelations. The challenges here are (i) modelling brain-wise correlation structure in a scalable manner and (ii) constructing clusterwise inference without relying on Gaussian Random Field Theory (GRFT) [20]. Our recent work provided a foundation for this goal by extending scan statistics and adopting permutation to obtain an empirical null distribution of the test statistic [28].

A summary of our methodological contributions is as follows:

- We use the nearest-neighbor Gaussian Process (NNGP) to get a close approximation of the true spatial autocorrelation structure with significantly reduced computation time.
- We propose a new cluster-enhanced statistic based on the *multivariate* test statistics leveraging an explicit spatial autocorrelation modelling.
- We use a *refitting-free* resampling method to compute the FWER-controlling threshold fast and accurately.

The proposed approach, called CLEAN (**C**lusterwise inference **le**veraging spatial **a**utocorrelations in **n**euroimaging) results in significantly improved sensitivity and specificity. We show that CLEAN controls FWER accurately through our permutation scheme and is also computationally highly efficient, which we will illustrate through a data analysis that took only a few minutes to complete.

The rest of the article is organized as follows. In Section 2, we describe the details of CLEAN and characterize its connections and differences to other clusterwise inference methods. In Section 3, we conduct extensive simulation studies to compare the performance of CLEAN to other methods, including the clusterwise inference without covariance modelling or the massive univariate analysis, with respect to different smoothing levels and SNRs. We then apply our method to a group of 44 subjects from the Human Connectome Project (HCP) who conducted the social cognition (theory of mind) task and the relational processing task twice (test-retest) [29]. We conclude with discussions in Section 4.

## 2. Methods

### 2.1. Data

We consider continuous (Gaussian) neuroimaging data collected in the spatial domain. Two examples are (i) activation in task-fMRI measured in cortical surface (exemplified by the Human Connectome Project) and (ii) cortical thickness or cortical area (structural MRI). The explicit pairwise distance information can be obtained on the cortical surface by computing the pairwise geodesic distance from the triangular mesh.

In this article, we use the two-level approach in analyzing task-fMRI. We obtain first-level contrast of parameter estimates (COPE) for each vertex from the fMRI time-series data for each subject, which we will treat as our ‘data’ for the group-level analysis [30].

### 2.2. Notations and model specifications

Let *i* = 1,…, *N* and *v* = 1,…, *V* be the indices for subjects and vertices of the triangulated brain surface. We assume that a location in a hemisphere of the brain is correlated only with locations within the same hemisphere. Let **z**_*i*_ be row nuisance covariate vectors (e.g. age or gender) and *x*_*i*_ be the covariate of interest (e.g. behavioral performance). The observed brain imaging data 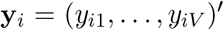 are modelled by **y**_*i*_ = **Z**_*i*_***α*** + **X**_*i*_***β*** + ***ϵ***_*i*_, where **Z**_*i*_ = **z**_*i*_ ⊗ **I**_*V*_, **X**_*i*_ = *x*_*i*_ ⊗ **I**_*V*_ and ⊗ is the Kronecker product. This is equivalent to the stack of the univariate (vertex-level) models but allows modelling the noise structure ***ϵ***_*i*_. For example, the one-sample test used in group-level activation in task-fMRI corresponds to *x*_*i*_ = 1 for all subjects while **Z**_*i*_***α*** is ignored. Similarly, the two-sample test for comparing group differences in means corresponds to **z**_*i*_ = 1 for all subjects and *x*_*i*_ = 1 if *i* is in group 1 and *x*_*i*_ = −1 if *i* is in group 2. Lastly, we assume 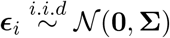 with 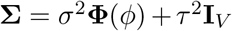, where **Φ**(*ϕ*) is the matrix of the spatial autocorrelation function (SACF) with a prespecified spatial kernel with a parameter *ϕ* and a pairwise distance matrix. For example, for the exponential kernel, 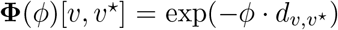 when 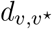 is the distance between vertices *v* and *v^*^*. *τ*^2^ is the nugget effect, the non-spatial variability of the data.

### 2.3. CLEAN

The key idea of CLEAN is three-fold. First, we model the spatial autocorrelation of the data and construct multivariate test statistics. Second, we construct an enhanced test statistic for each vertex by borrowing information from the spatial domain. Third, we use resampling to estimate the variance of each statistic and to control for the family-wise error rate (FWER).

#### 2.3.1. Multivariate test statistics

Our null hypothesis is *H*_0_: ***β*** = **0**, with a specific goal of identifying nonzero indices of ***β***. Instead of obtaining 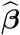 and its covariance matrix, we propose using the score test statistics, which can be obtained by fitting the model assuming that *H*_0_: ***β*** = **0** is true (i.e. the null model is **y**_*i*_ = **Z**_*i*_***α*** + ***ϵ***_*i*_). The score test is equivalent to the Wald test with large sample size, but the score test has a dramatically lower computational cost when using permutations [28]. The multivariate score statistic for testing *H*_0_ is

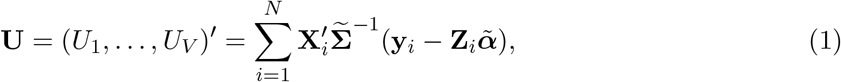

with 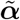 and 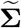 obtained by fitting the null model. We propose using a one-step update using the generalized least squares (GLS) framework to reduce the computational burden. Specifically, we regress {**y**_*i*_} on {**Z**_*i*_} via ordinary least squares (OLS) and obtain residuals, and use them to obtain 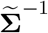] and then to update 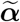. Using simulations and a real data analysis, we show that the one-step update sufficiently controls for false positives and preserves high statistical power.

#### 2.3.2. Spatial autocorrelation modelling using the nearest-neighbor Gaussian process (NNGP)

A key challenge to constructing the test statistics in Equation (1) is inverting 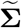. Due to the large number of vertices, it is computationally infeasible to invert this matrix without using an approximation. To resolve this problem, we use the nearest-neighbor GP (NNGP) to obtain an accurate approximation of the true spatial Gaussian process and alleviate the computational cost [31]. The NNGP assumes that the Cholesky decomposition of **Σ**^−1^ consists of sparse matrices and uses nearest neighbors to approximate the true GP [31, 32]. Specifically, we replace **Σ**^−1^ with 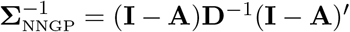 in Equation (1), where **D** is a diagonal matrix and **I** − **A** is a *sparse* lower triangular matrix constructed by *J* predefined nearest neighbors and {*σ*^2^, *τ*^2^, *ϕ*} (see Figure 1). A detailed procedure of obtaining **A** and **D** is illustrated in the Appendix A. *J* balances the quality of the approximation of true Gaussian process and computational efficiency. Following the suggestions by Datta et al. [31], we set *J* = 50, but one can flexibly change this based on the resolution of the images. For example, when using the exponential kernel, one can first determine the smallest *d* such that the parametric correlation 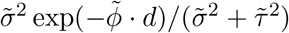 is less than a certain threshold (e.g. 0.05) and use the number of vertices within the neighbor of radius *d* as *J*. As shown in Appendix C, setting *J* = 50 yielded qualitatively similar results to *J* = 100 in the data analysis. Because obtaining 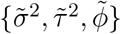 would be relatively easy by using a method-of-moment approach, as outlined in the Appendix A2, obtaining 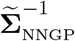 as a significantly reduced computational cost 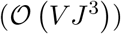 than naively inverting 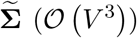. In fact, the computational burden only increases *linearly* with *V* [31], which is very practical since the number of vertices is often very large in neuroimaging. Also, fitting an NNGP can be further accelerated using parallel computing [32].

**Figure 1:**
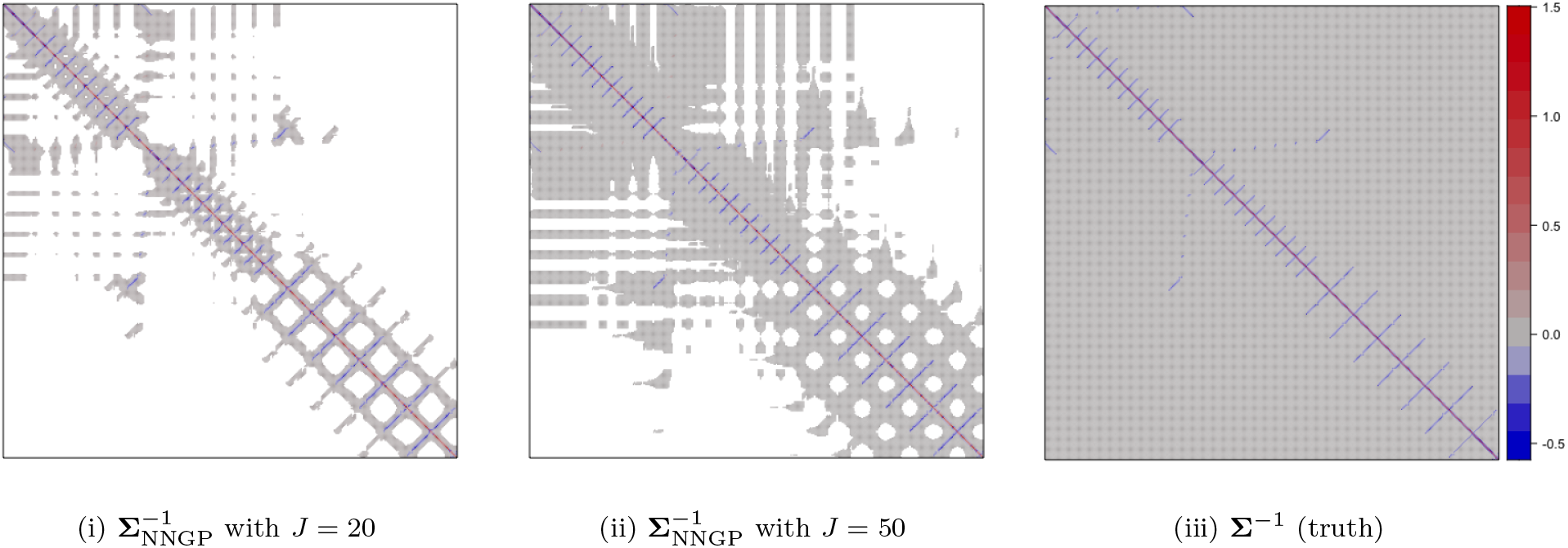
Illustration of NNGP, using a 20 *×* 20 rectangular grid (*V* = 400), using the exponential covariance function with *σ*^2^ = 2, *τ*^2^ = 0.5, *ϕ* = 0.5. The true **Σ**^−1^ as well as the NNGP approximations with different neighbor sizes (*J* = 20, 50) are shown. White entries in (i) and (ii) correspond to entries that are exactly 0.

Unlike other approximation methods such as the spatial Gaussian predictive processes (SGPP), the NNGP is a well-defined GP derived from the actual spatial GP and uses the same parametrizations, with fairly accurate approximations [31]. Since the NNGP is not uniquely defined and depends on the initial vertex that constructs the process, we choose the initial vertex where the sum of the pairwise distances is the smallest. However, as argued by Datta et al. [31] and as shown in Appendix C, the results are robust to the choice of the initial vertex.

With a minimal smoothing level, we find that the exponential covariance structure closely reflects the empirical covariance computed using data on the spherical surface, which agreed with Risk et al. [12] (Appendix D). We estimate {*σ*^2^, *τ*^2^, *ϕ*} by minimizing the sum of squared differences between empirical and theoretical spatial covariances. The detailed estimation procedure is provided in the Appendix B.

#### 2.3.3. Clusterwise inference

Spatial modelling would need to model both the mean structure and the error structures to achieve higher power [4, 12]. Using the score test statistics (Equation (1)), we propose to use clusterwise inference to model the mean structure of the data by using neighboring vertices’ test statistics. We first define *N*_*r*_ (*v*) be the collection of vertices whose distances from vertex *v* is less than *r*. The cluster-enhanced test statistic for vertex *v* is defined by *T*(*v*) = max{*T*_*r*_ (*v*)|*r* ∈ Γ} where *T*_*r*_ (*v*) is a standardized average of the score test statistics within *N*_*r*_ (*v*),

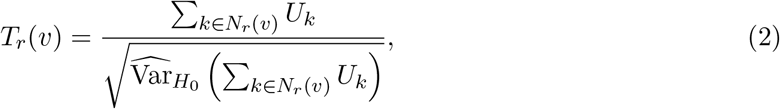

and Γ is a set of radii, with each element corresponding to the set of neighbors *N*_*r*_ (*v*).

When *H*_0_ is true, the distribution of **U** is multivariate normal with zero mean [28, 33]. Therefore, the distribution of *T*_*r*_ (*v*) is approximately standard normal for any *r* and *v*. If *N*_*r*_ (*v*) specifies the true signal regions, *T*_*r*_ (*v*) will be more powerful in rejecting *H*_0_ than univariate inference. For illustration, Figure 2 describes three candidate clusters such that the true signal region is (i) correctly specified (ii) underspecified, and (iii) overspecified. The ideal goal is to have correctly specified clusters as shown in (i), since under/over-specification of the clusters can affect statistical power. Since the true signal location is unknown, *T*(*v*) uses a combination of candidate clusters with various sizes to determine the location of the signal locations in a data-adaptive manner.

**Figure 2:**
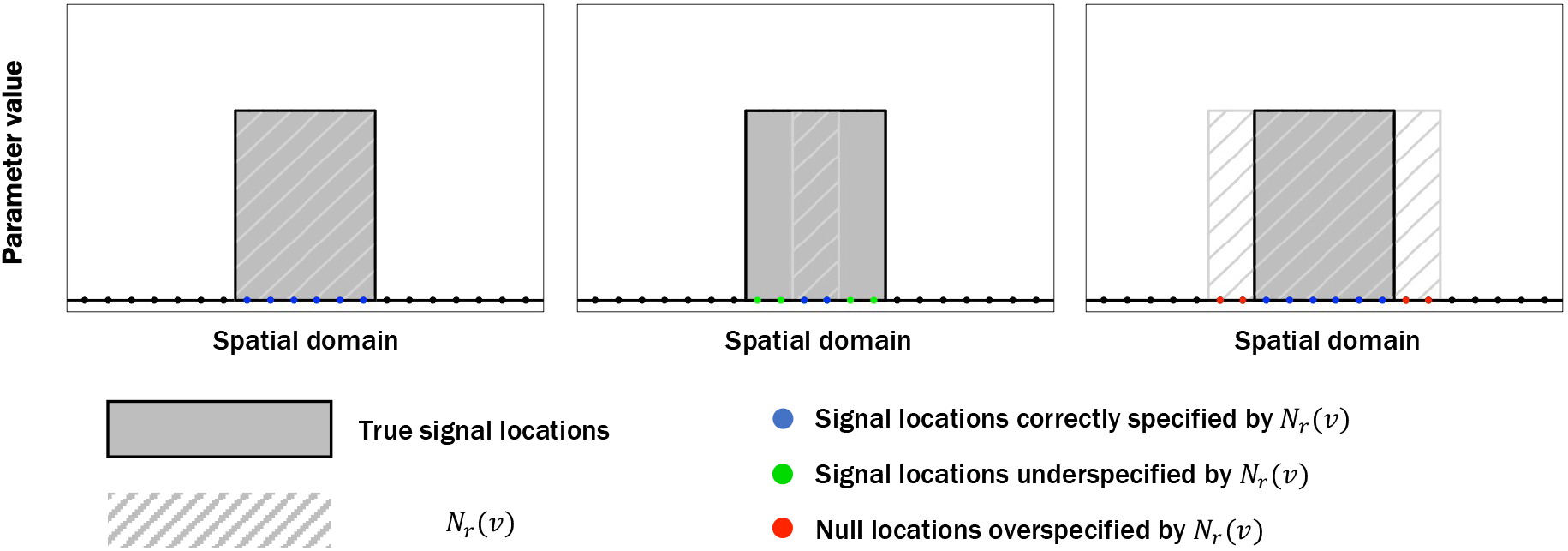
Illustration of possible cases regarding the choice of candidate clusters. (Left) Correctly-specified candidate cluster. (Middle) Under-specified candidate cluster, which does not include all signal locations. (Right) Over-specified candidate cluster, which contains null vertices.

**Figure 3:**
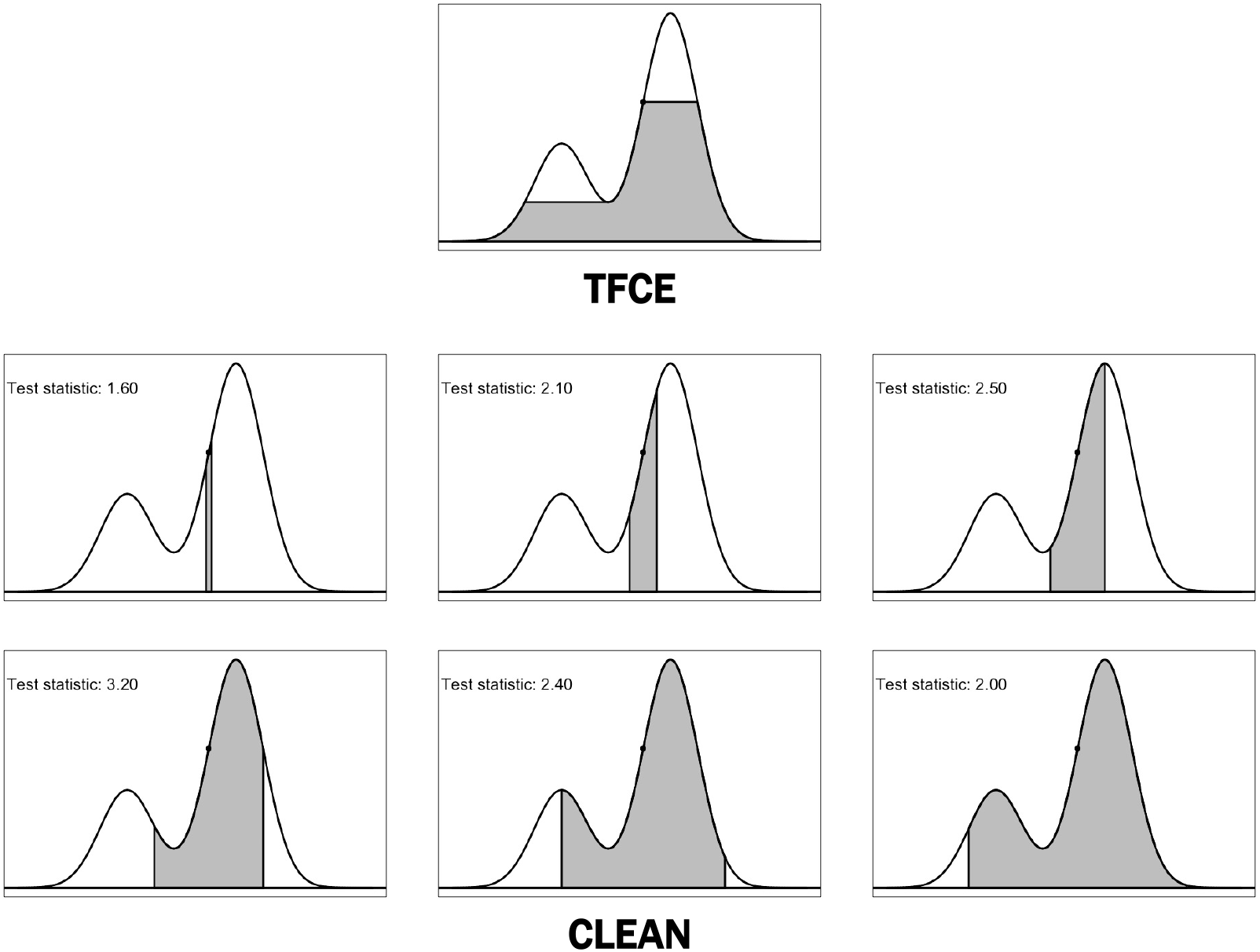
A conceptual illustration of the difference between TFCE and CLEAN. The *y* axis denotes for the test statistic (either univariate or multivariate). TFCE uses the *distribution* of the test statistics to obtain enhanced test statistic, defined by *enhanced area*. The support depends on the distribution of the test statistics and TFCE statistic does not follow an explicit probability distribution. CLEAN considers a fixed yet wide range of supports predefined by the neighbor information and takes the maximum of the *test statistics* as the enhanced statistic for the voxel. The covariance of the test statistics is used to compute a test statistic in each domain, and each test statistic follows 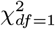 asymptotically (in two-sided tests). In this example, a cluster with test statistic 3.20 is preferred to other candidates, when all these test statistics exceed the FWER-controlling threshold.

#### 2.3.4. Refitting-free resampling

Nonparametric approaches are necessary to control false positives accurately in clusterwise inference [20]. We use a resampling approach to (i) obtain the empirical distribution of *T*(*v*) under the null hypothesis *H*_0_ and (ii) set a threshold that controls FWER. Among existing approaches [34], we use the method of Draper and Stoneman [35] which was also proposed in our early work with the score test framework [28]. In particular, by combining the score test with the proposed resampling method, obtaining the permuted test statistic only requires matrix multiplications (applying Equation (1) with resampled **X**_*i*_). This approach is computationally highly efficient since it does not require refitting the full model.

In the one-sample test, we flip signs of each individual’s image and construct permuted score statistics. In the two-sample test, we permute *x*_*i*_ across subjects. We then use the sample covariance of the sign-flipped/permuted score statistics to obtain the empirical null distribution of *T*(*v*) (i.e. the denominator of Equation (2)). The permuted score statistics can also be used to set a threshold that controls family-wise error rate (e.g. the (1 − *α*)th quantile of the maximum of the permuted test statistics).

We note that, because the denominator of Equation (2) is obtained by the sample covariance of the permuted score statistics, having a sufficient number of permutations is necessary, especially when |*N*_*r*_ (*v*)| is large. Therefore, we recommend having at least 10,000 resamples to robustly estimate the empirical distribution of *T*_*r*_ (*v*) in Equation (2).

#### 2.3.5. Remarks

- *On choosing the neighbor set* Γ: The performance of the proposed method depends on the specification of neighbors *N*_*r*_ (*v*). While a large radius size *r* would result in a higher power when the actual size of a cluster is large. However, every clusterwise inference that borrows information from the neighbors suffers from the loss of specificity. Therefore, an appropriate level of radii is necessary to balance power/sensitivity and specificity. Therefore, we use Γ = {1mm, 2mm,…, 20mm}. However, the elements of Γ can be set by investigators a priori. Defining neighbor sets by distances provides a uniform degree of combination in any resolution of the image. Our earlier work used the number of nearest neighbors to specify candidate sets [28], which is another possible way. Still, it does not give uniform candidate sets with different resolutions of the image.
- *On the nugget effect τ*^2^: While existing methods modelling spatial autocorrelation ignored or removed *τ*^2^, our data analysis reveals that non-ignorable non-spatial variation is present unless data has been sufficiently smoothed. Therefore, ignoring the nugget effect would result in inaccurate control of FWER and potentially affect statistical power.
- *Comparisons to existing clusterwise inference methods*: The proposed method adaptively combines test statistics across predefined neighbors, including locations with negative test statistics. Existing methods, such as TFCE, computes cluster-enhanced test statistic for each location by aggregating test statistics in the spatially connected region (i.e. locations with non-negative test statistics). An assumption on the smoothness is a key idea that increases sensitivity in TFCE. Motivated partially by Gaussian Random Field Theory, *E* = 0.5 and *H* = 2 is used to set the TFCE statistic as follows:

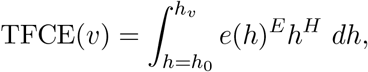

where *h*_*v*_ is the univariate test statsitic (*z* score) for location *v* and *e*(*h*) is the ‘support’ (i.e. area of the locations) corresponding to statistic *h* [14]. CLEAN is better motivated when the shape of test statistics in the spatial domain is not smooth. It is useful in our setting where spatial smoothness (i.e. autocorrelation) is *modelled* directly. By restricting the maximum of Γ, it also limits the degree in the spatial domain that test statistics borrow information. As a result, it avoids the issue that the clusterwise inference method is prone to low specificity when the cluster size is large [16]. CLEAN also has some overlaps with LISA (Local Indicators of Spatial Association) [15] in that the cluster-enhanced statistic is computed by aggregating test statistics in pre-defined neighbor sets. Given *z*_*v*_, the *z* score for voxel *v*, LISA uses the following statistic:

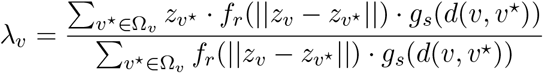

where Ω_*v*_ is a neighbor set (analogous to *N*_*r*_ (*v*)) and *f*_*r*_ (·) and *g_s_*(·) are kernel functions for differences in *z* scores and pairwise distance, respectively. LISA applies the mixture model framework and uses permutation to control false discovery rate (FDR), whereas CLEAN controls FWER. Also, we allow multiple neighbor sets for each *v* to reflect the true and unknown size of signal clusters data-adaptively. Also, CLEAN models spatial autocorrelations explicitly, which is not the case in LISA.
- *Software*: CLEAN is currently available at https://github.com/junjypark/CLEAN as an `R` package. It uses `C++` to boost computational efficiency, and additional gain is feasible by using the parallel computing supported by the package. Without parallel computing, applying CLEAN to a neuroimaging data of 10,000 vertices and 44 subjects took less than 5 minutes using 10,000 permutations on a Macbook Pro with 2.3 GHz Quad-Core Intel Core i5 and 16GB RAM.

## 3. Data analysis

### 3.1. Data Collection

We obtained data from the Human Connectome Project (1200 Subjects Data Release) who completed the social cognition (SC) task and the relational processing (RP) task and passed quality control. It resulted in 1050 subjects for SC and 1041 subjects for RP. In SC, we considered the Theory of mind (TOM) versus Random contrast. In RP, we considered Relational vs Match contrast. As previous analyses revealed that the effects in SC are strong and widely spread over the brain and the effects in RP are sparse and weak, these tasks are appropriate to see how each method performs with respect to different effect sizes. For analysis, we chose 44 subjects who completed the task twice (test-retest). These subjects included 31 females and the age ranged between 22 and 35.

The fMRI data were processed with HCP’s minimal surface processing pipeline. Using the fMRI time series with 2mm surface smoothing, we obtained the contrast of parameter estimates (COPEs) from each subject. To reduce the dimension of the data, we resampled 10K vertices per hemisphere using the R package `ciftiTools` [7], yielding 9394 and 9398 vertices in the left and right hemispheres, respectively.

Our analysis primarily focuses on 44 subjects who completed the task twice. Because of the large number of samples used in this study, we pooled information across all subjects (more than a thousand) to obtain an image of effect sizes to evaluate the validity of the proposed method. We computed Cohen’s *d* for each vertex as a measure of the effect size, given by the absolute value of the average of the COPEs divided by the standard deviation of COPEs across all subjects. Following Geuter et al. [36] and Cohen [37], we thresholded the effect size image using 0.2, 0.5, and 0.8 (see Figure 4).

**Figure 4:**
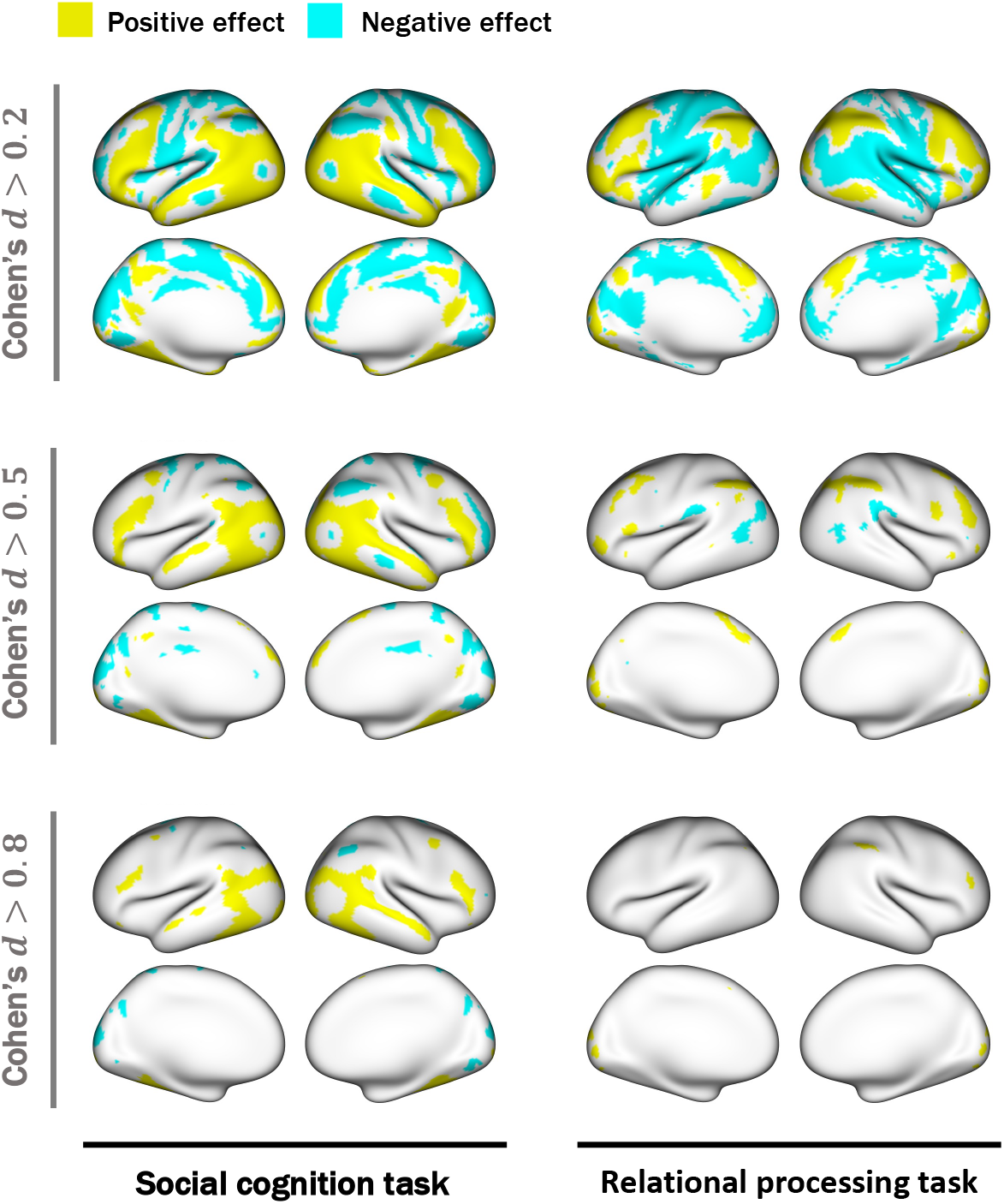
The effect size image (lateral and medial views) of the HCP’s social cognition task with thresholds 0.2, 0.5, and 0.8. The effect sizes were computed using the COPE information from 1050 (social cognition) and 1041 (relational processing) subjects. Signs are preserved for visualizations. The figures are displayed on the inflated surface.

### 3.2. Results

We fitted three different models to analyze 44 subjects’ test data: (i) CLEAN, which leverages the spatial autocorrelation in clusterwise inference, (ii) clusterwise inference without leveraging the spatial autocorrelation, and (iii) massive-univariate analysis. Approach (ii) is equivalent to CLEAN but without using NNGP. All methods used permutation to control FWE at the rate of 0.05.

The areas of significance are illustrated in Figure 5. In both tasks, CLEAN was superior to the other approaches in capturing areas of small effect sizes and small signal clusters. Specifically, CLEAN identified most areas with Cohen’s *d* of 0.5 or greater. It also identified a few areas with Cohen’s *d* less than 0.5. Massive univariate analysis identified areas with high effect sizes but underperformed when the effect size is small. We also note that the CLEAN without spatial autocorrelation modelling (Approach (ii)) resulted in smoother areas of significance when compared to the massive univariate analysis. Because in clusterwise inference one may only conclude that ‘at least one location’ within the cluster is truly a signal location, Approaches (ii) and (iii) were qualitatively similar. Moreover, Approach (ii) oversmoothed the areas and reported a false direction of the effect in (ROI), which implies that it could be prone to low specificity.

**Figure 5:**
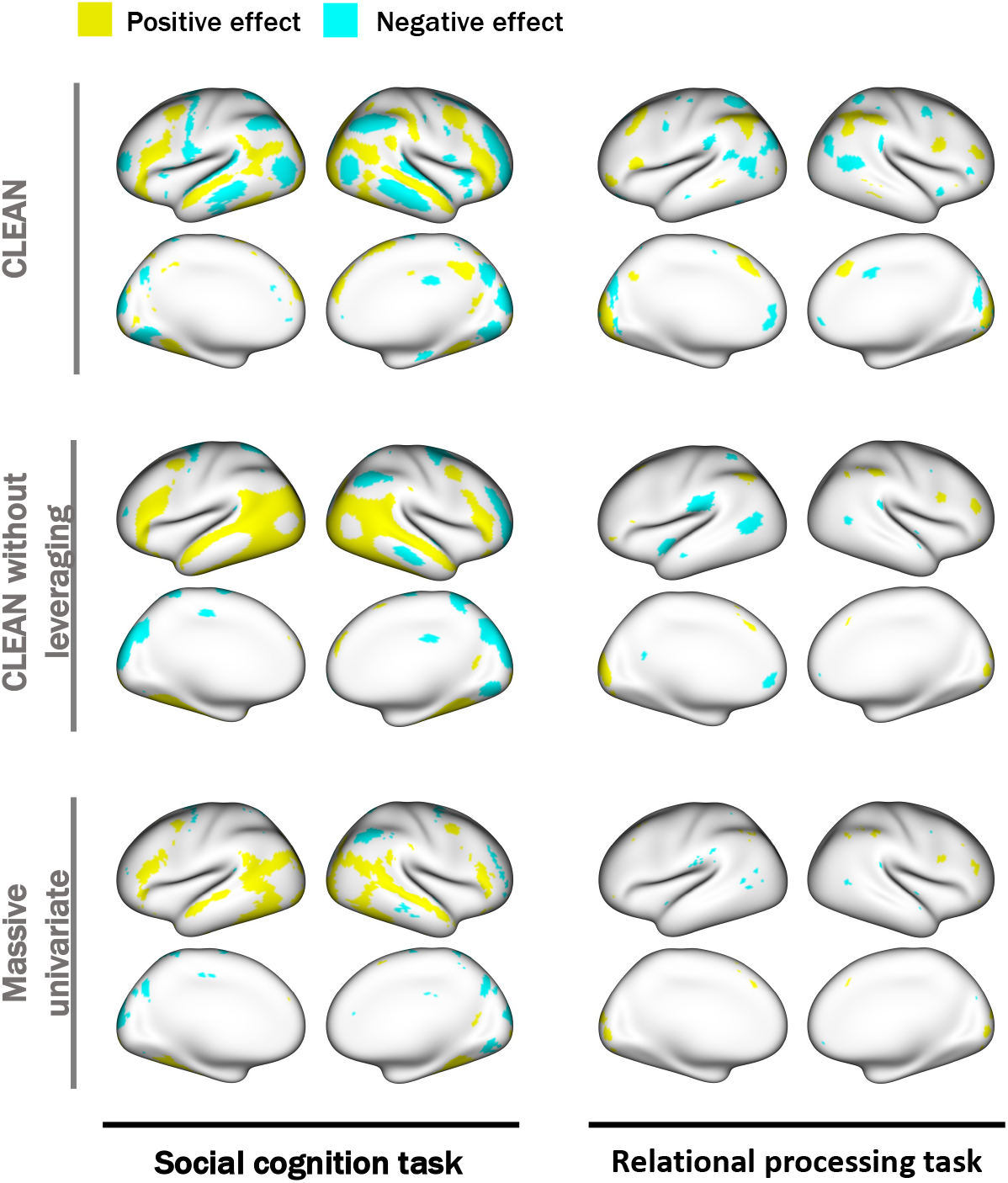
Analysis of 44 subjects who completed the tasks twice (test-retest) in the HCP. FWER is controlled at the rate of 0.05. CLEAN identified more regions that aligned with population-level effect size images in Figure 4.

### 3.3. Analysis of the retest data

We also analyzed the retest data from the same set of subjects. The result is shown in Figure 11 in Appendix E. Overall, there is a high agreement on the detected signal locations between the test and retest data, supporting the reproducibility of the results using CLEAN. However, we do not compare different methods based on the empirical agreement measure (e.g. proportion of overlap) because the interpretation of clusterwise inference cannot be made on a vertex level. We therefore conducted empirical simulations in Section 3.5.3 to evaluate vertex-wise reproducibility.

### 3.4. Sensitivity analysis

As discussed in Section 2, NNGP is not uniquely defined and its specification depends on the initial vertex as well as the number of neighbors used to specify the process. We evaluated the sensitivity of the result by trying 10 randomly chosen initial vertices to construct NNGP with *J* = 50 and 100 neighbors. As shown in Figure 9 of Appendix C, there is a high agreement among the results with different initial values when *J* is fixed. It supports the robustness of the proposed method. The results using *J* = 50 and *J* = 100 were qualitatively similar, which agrees with Datta et al. [31].

### 3.5. Simulation studies

#### 3.5.1. Family-wise error rate

We evaluated empirical FWER using two designs: inference for the mean (one-sample test) and inference for the difference (two-sample test). In both designs, each simulated data consisted of *N* = 30 left-hemisphere images from the HCP’s social cognition task. In the one-sample test, we generated noise-only images by first computing the “population-mean” image by taking the average of all 1050 subjects’ COPE images, then subtracting the population-mean image from each subject’s COPE image. In the two-sample test, we used the original data, but we randomly assigned subjects into two groups with 15 subjects each. Because of the random assignment, it is expected that the difference of the means of the two groups has a zero mean. Therefore it is valid to evaluate FWER. Using the FWER of 0.05, we computed the empirical FWER, the proportion of datasets that at least one vertices were declared as signals.

Each simulation was repeated 1000 times. The empirical FWERs were 0.051 (one-sample test) and 0.048 (two-sample test), respectively, suggesting that the proposed method accurately controls the false positives.

#### 3.5.2. Power

We evaluated statistical power of the global hypothesis using simulations. We first defined the “signal-only” image as the “population-mean” image where the effect size is greater than 0.8. We then scaled signal-only image to have the range between 0.01 and 5. For each subject’s noise-only image, we added the signal-only image scaled by *γ* = 1, 2, · · ·, 10. For each *γ*, the power was defined by the proportion of simulated datasets (out of 1000) rejected by each method, using the FWER of 0.05. We first verified that the competitors controlled FWER at the nominal level 5%.

The power curve for each method is summarized in Figure 6. The power curve for CLEAN is uniformly higher than the curves of the competitors. The simulation results reveal that CLEAN without spatial autocorrelation modelling is equivalent to massive univariate analysis in terms of power. Notably, for the competitors, 80% power is obtained when *γ* = 6, whereas CLEAN achieves the same power with lower effect size (*γ* ≈ 5.5).

**Figure 6:**
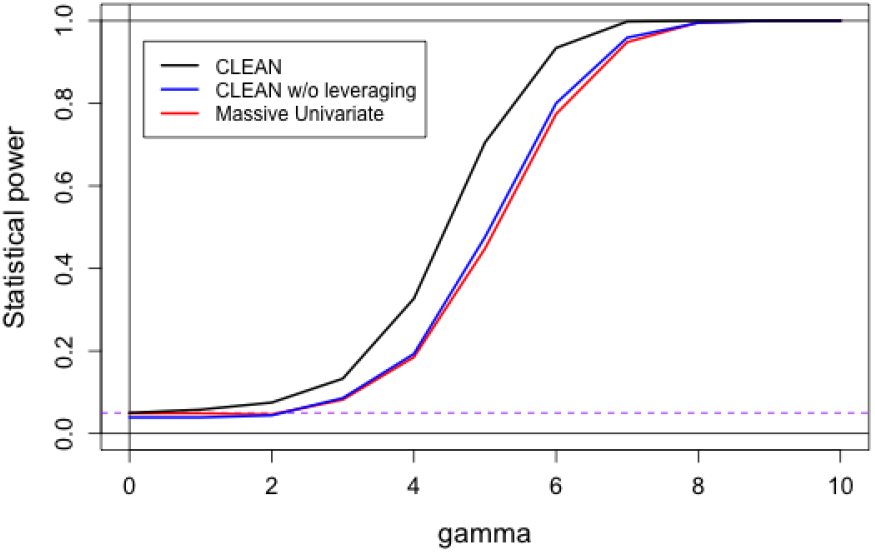
Power analysis from the simulation studies. Simulated data is generated by adding signal images (thresholded by high effect size) to the noise-only images from the HCP social cognition task. Purple dotted line is the FWER (0.05).

#### 3.5.3. Vertex-level reproducibility

Note that power is insufficient to evaluate how each method identified signal regions because rejecting the null hypothesis only yields that ‘at least one’ vertex is statistically significant. Therefore, we used the SC and RP data from HCP to evaluate the reproducibility of scientific findings.

In this section, we used varying numbers of sample sizes: *N* = 20, 30, 40. For each fixed sample size, we generated 500 simulated datasets of the COPE images from each task of the HCP. Then we fitted each method and extracted the detected signal vertices for each simulated data. We visualized these differences in Figure 7 by computing the proportion of times selected by each method for every vertex (out of 500 simulated datasets). The proposed approach still outperformed the others in detecting new signal regions with higher reproducibility. Notably, when *N* = 40, 16.7% (SC) and 4.3% (RP) of the vertices had over 80% reproducibility with CLEAN, which were substantially higher than CLEAN without leveraging (14.9% and 0.8%) and the massive univariate analysis (5.4% and 0.4%). Even when we used the 50% cutoff for the reproducibility, CLEAN (24.8% and 9.1%) is superior to CLEAN without leveraging (21.6% and 3.3%) and the massive univariate analysis (10.2% and 0.6%). Also, we observe in Figure 7 that CLEAN successfully identifies new localized regions with high reproducibility, while CLEAN without leveraging tended to smooth wider, which agrees with Figure 5.

**Figure 7:**
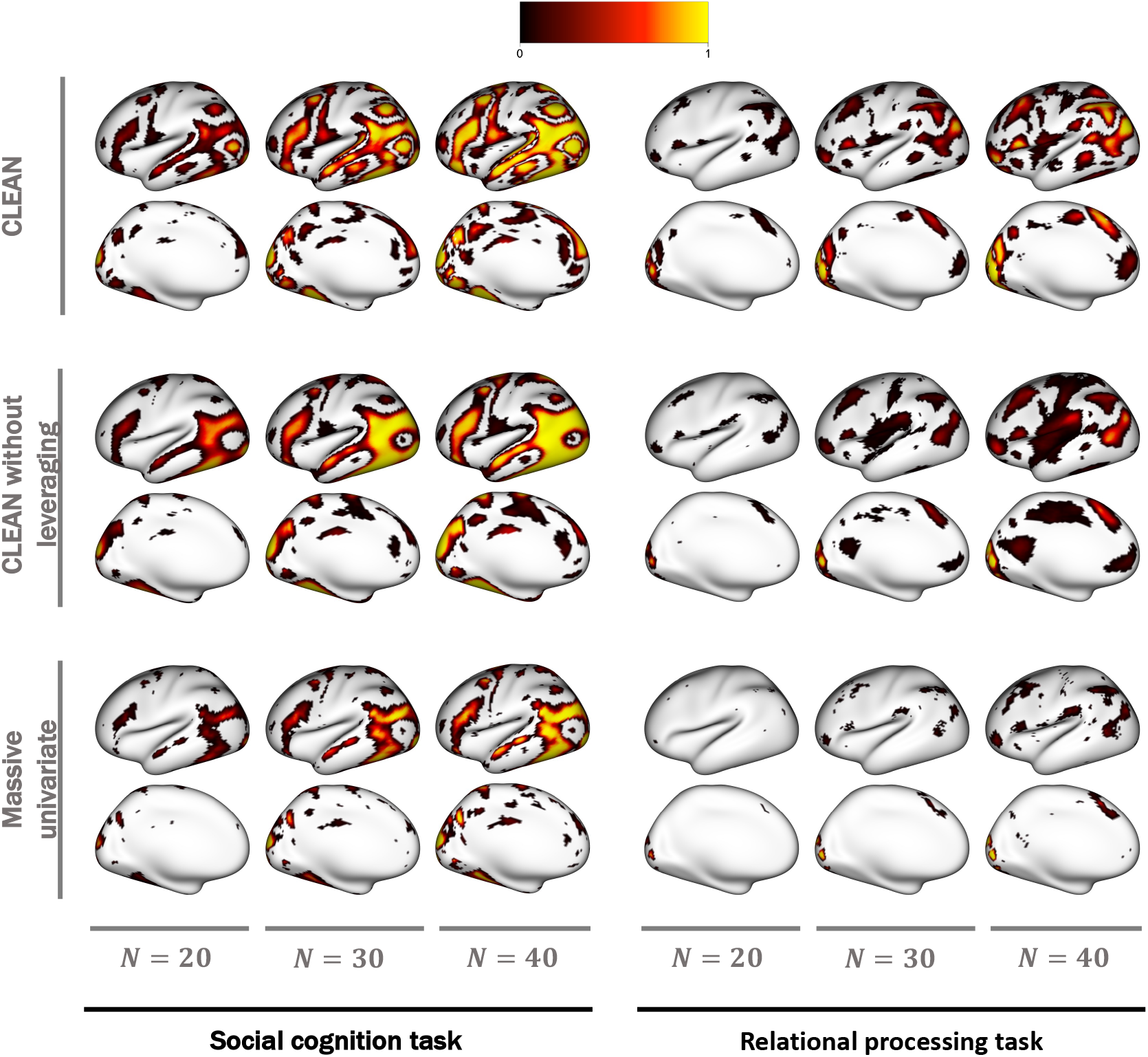
Vertex-level reproducibility for each method. 500 simulated data with different sample sizes (*N* = 20, 30, 40) were generated. For each vertex, the proportion of time detected by each method is displayed.

We also defined three different signal images based on the effect size (Cohen’s *d* greater than 0.8, 0.5 and 0.2), and computed the sensitivity. In each dataset, we computed the number of overlapping vertices between the signal image and identified vertices divided by the total number of signal vertices. We then averaged the proportion across 500 simulated datasets. Figure 8 shows that CLEAN, regardless of the modelling of spatial autocorrelations, yielded higher sensitivity than massive univariate analysis. In SC, CLEAN without autocorrelation modelling yielded slightly higher reproducibility than the proposed approach with medium and large effect sizes, but CLEAN slightly better when effect size is small. It is understandable because the brain activation in SC is global and widespread, and CLEAN without leveraging spatial autocorrelation tended to oversmooth the areas of significance. In RP, where effect size is small and more localized, CLEAN without leveraging did not gain much by oversmoothing. Therefore, it is understandable that CLEAN resulted in higher sensitivity in RP than the competitors.

**Figure 8:**
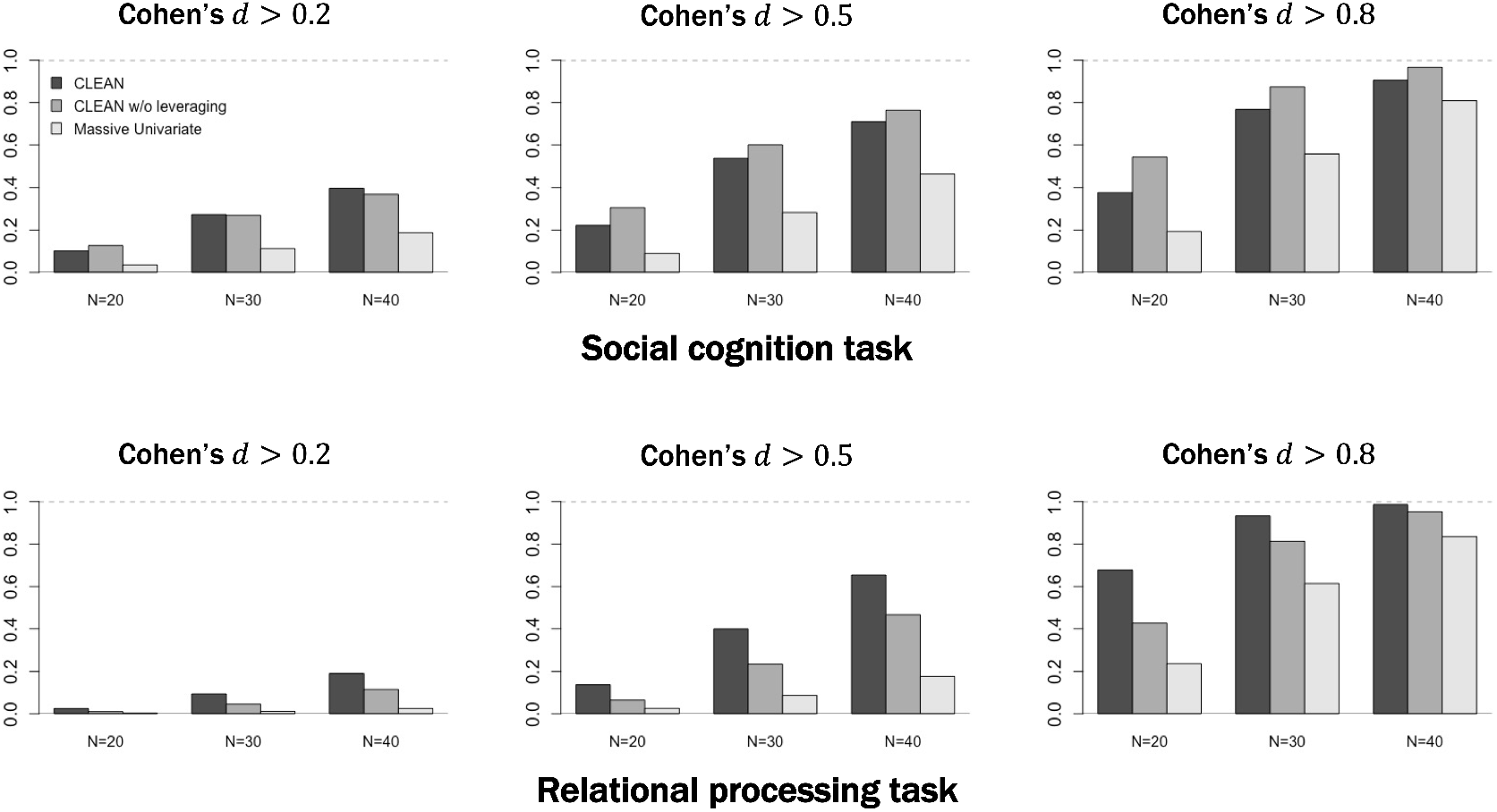
Sensitivity with respect to different ‘signal’ images binarized by different Cohen’s *d* levels. We averaged the sensitivity across 200 simulated datasets. In the social cognition task where activation levels are strong and widespread, CLEAN without spatial autocorrelation modelling tended to have higher sensitivity by oversmoothing the activation regions. In the relational processing task where activations levels are weak and localized, CLEAN yielded higher sensitivity.

#### 3.5.4. Summary

Benefiting from the large database of the Human Connectome Project, we validated that CLEAN controlled false positives accurately, which is attributed to the use of permutations. CLEAN also showed higher power and vertex-wise reproducibility that agreed closely on the population-level effect sizes. Comparing it to two competitors (massive univariate analysis and CLEAN without leveraging), we showed that both spatial autocorrelation modelling and cluster-wise inference contributed to the higher reproducibility.

## 4. Discussion

We propose a novel clusterwise inference method for neuroimaging data called CLEAN that models the spatial autocorrelation explicitly through a spatial Gaussian process for improved sensitivity and specificity. By analyzing the HCP data and using empirical simulations, we demonstrated that CLEAN controls FWER accurately and outperforms massive univariate analysis and clusterwise inference without modelling spatial autocorrelations. CLEAN is computationally very efficient and applicable to a wide range of neuroimaging data with spatial autocorrelation. Our method is publicly available as a form of an R package at https://github.com/junjypark/CLEAN, and it supports parallel computing to further reduce the computational cost.

The key ideas that played a central role in the proposed approach are the nearest-neighbor Gaussian process (NNGP), the refitting-free permutation via the score test framework, and the construction of the cluster-enhanced test statistic. NNGP provided a close approximation to the true spatial Gaussian process that captured most spatial variations of the noise. This is very useful when working with images with higher resolutions because the construction of NNGP can be parallelized, and the computational cost increases linearly with respect to the number of vertices. Also, our method is based on the score test, which does not require refitting a multivariate model multiple times. Moreover, the proposed clusterwise inference procedure constructs test statistics based on prespecified neighbor sets and avoids restrictive GRFT. Finally, quantifying uncertainty, or a degree of the spatial extent that benefits from the clusterwise inference, is straightforward.

The proposed method addresses a few issues on existing clusterwise inference. First, existing methods have used a degree of Gaussian smoothing to increase the signal-to-noise ratio, but we point out that it makes the null location into the pseudo-signal location. Because, in theory, a spatial location is either a null or a signal location, identifying a pseudo-signal location as a signal location would only decrease specificity. Our approach does not rely on a wide extent of smoothing because it models spatial autocorrelation explicitly and in a scalable manner. Second, Woo et al. [16] discussed that clusterwise inference is prone to low specificity when the actual cluster size is large. This is a common problem in clusterwise inference, and one may only conclude that *at least one* vertex within the declared regions is significant. Although the proposed method also falls in this category, one can control the maximum distance where information is borrowed. This is contrary to other methods where the cluster-enhanced statistic is constructed using non-negative statistics of the neighboring locations. The cluster-enhanced statistic is also straightforward in meaning, and it determines how well ‘collapsing’ data across the cluster maximizes the mean (1-sample test), the difference in means (2-sample test), correlation (simple linear regression) or partial correlation (GLM).

The proposed approach has room for improvement. First, our analysis focused on brain imaging data, including activation in task-fMRI, measured on the cortical surface that shows an explicit spatial autocorrelation. Extending it to 3D (volumetric) fMRI data would be straightforward as long as the assumption on the stationarity of the spatial autocorrelation is valid. We leave it as future work. Second, our approach is currently restricted to group-level analyses (i.e. population-level inference). A method for analyzing a single subject’s fMRI would be an interesting extension of the current work. Third, we used the spherical surface to compute the geodesic distance matrix to ensure that subjects are registered onto the same surface [12, 23]. However, this could be sensitive to the distortion of the distance information, and other surfaces, such as the midthickness surface, might be more appropriate to reflect the cortex. Fourth, the proposed model assumes a stationary Gaussian process, meaning that any pair of vertices would reveal the same degree of autocorrelation if the distances are the same. Although the proposed method controlled FWE accurately, accounting for potential nonstationarity would further improve statistical power.

## 5. Conclusions

We proposed a novel and general statistical method that simultaneously models both the mean structure and the spatial covariance structure. It is a powerful method that successfully unveiled regions with low-medium effect sizes. Our method is a general approach applicable to many neuroimaging data with spatial autocorrelations. It is computationally highly efficient and is implemented as a R package.

## Acknowledgements

Data were provided by the Human Connectome Project, WU-Minn Consortium (Principal Investigators: David Van Essen and Kamil Ugurbil; 1U54MH091657) funded by the 16 NIH Institutes and Centers that support the NIH Blueprint for Neuroscience Research; and by the McDonnell Center for Systems Neuroscience at Washington University. We acknowledge the support of the Data Sciences Institute at the University of Toronto.

## Appendix

## A. Construction of the nearest-neighbor Gaussian process (NNGP)

Provided a *V* × *V* pairwise distance matrix 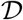 and spatial covariance parameters {*σ*^2^, *τ*^2^, *ϕ*}, constructing NNGP with *J* neighbors is summarized by two steps: (i) Constructing nearest-neighbor information and (ii) using it to compute 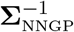.

The step (i) constructs the neighborhood information. Let 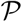 be a *V* × *J* matrix and let 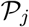 the *j*th row of 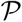. We first choose an initializing vertex *v*\* and store it to the element of 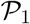. Then, for rows *j* > 1, 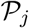 include a new vertex *v* closest to the vertex chosen in the row *j* − 1 and min(*j* − 1, *J* − 1) closest vertices of *v* chosen among the collection of vertices of 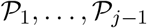.

The step (ii) is summarized by the following steps.

1. Let **A** = **0**_*V*×*V*_ and **D** = **I**_*V*_, and let **D**[*v*\*, *v*\*] = *σ*^2^ + *τ*^2^.
2. For *j* = 2,…, *V*,

a. Construct 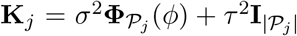, where 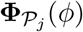 is a submatrix of **Φ**(*ϕ*) constructed by the indices 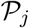.
b. Let *k* be the last element of 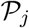 and 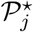 is the remaining elements, with the order preserved.

- 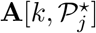 is the vector **x** that solves the linear system 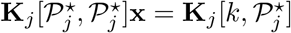.
- 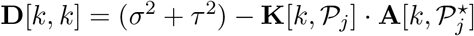 Note that this step can be easily parallelized.
3. The resulting 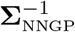 is 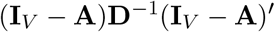

## B. Estimation of the spatial covariance parameters

We propose estimating spatial covariance parameters via the covariance regression analysis proposed by Zou et al. [38]. With *ϕ* provided, a consistent estimator of {*σ*^2^, *τ*^2^} can be obtained by minimizing

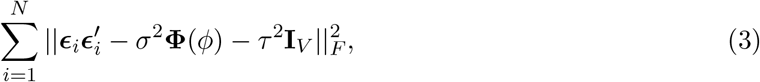

where 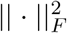 is the squared Frobenius norm of a matrix (sum of squared elements). With some algebra, a closed-form solution for these parameters is provided by

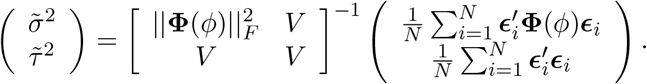

Therefore, it is sufficient to find *ϕ* to estimate {*σ*^2^, *τ*^2^} in which {*σ*^2^, *τ*^2^, *ϕ*} conditionally minimizes Equation (3), where a nonlinear optimization can be used. We the use 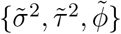 as resulting parameter estimates. When there are *q* covariates used to extract ***ϵ***_*i*_ including the intercept, then we multiply 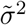 and 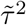 by *N*/(*N* − *q*) to adjust for the degrees of freedom. In Appendix F, we use simulations to show that the proposed loss function successfully recovers the true parameter estimates on average.

## C. Sensitivity analysis of the NNGP

We used 44 subjects from the HCP’s social cognition task, who completed the task twice (test-retest). Fixing the number of nearest neighbors *J* for specification of the NNGP, we used 5 different vertices that initialize the process. The results shown in Figure 9 reveal that the proposed method is robust to the initial vertex.

**Figure 9:**
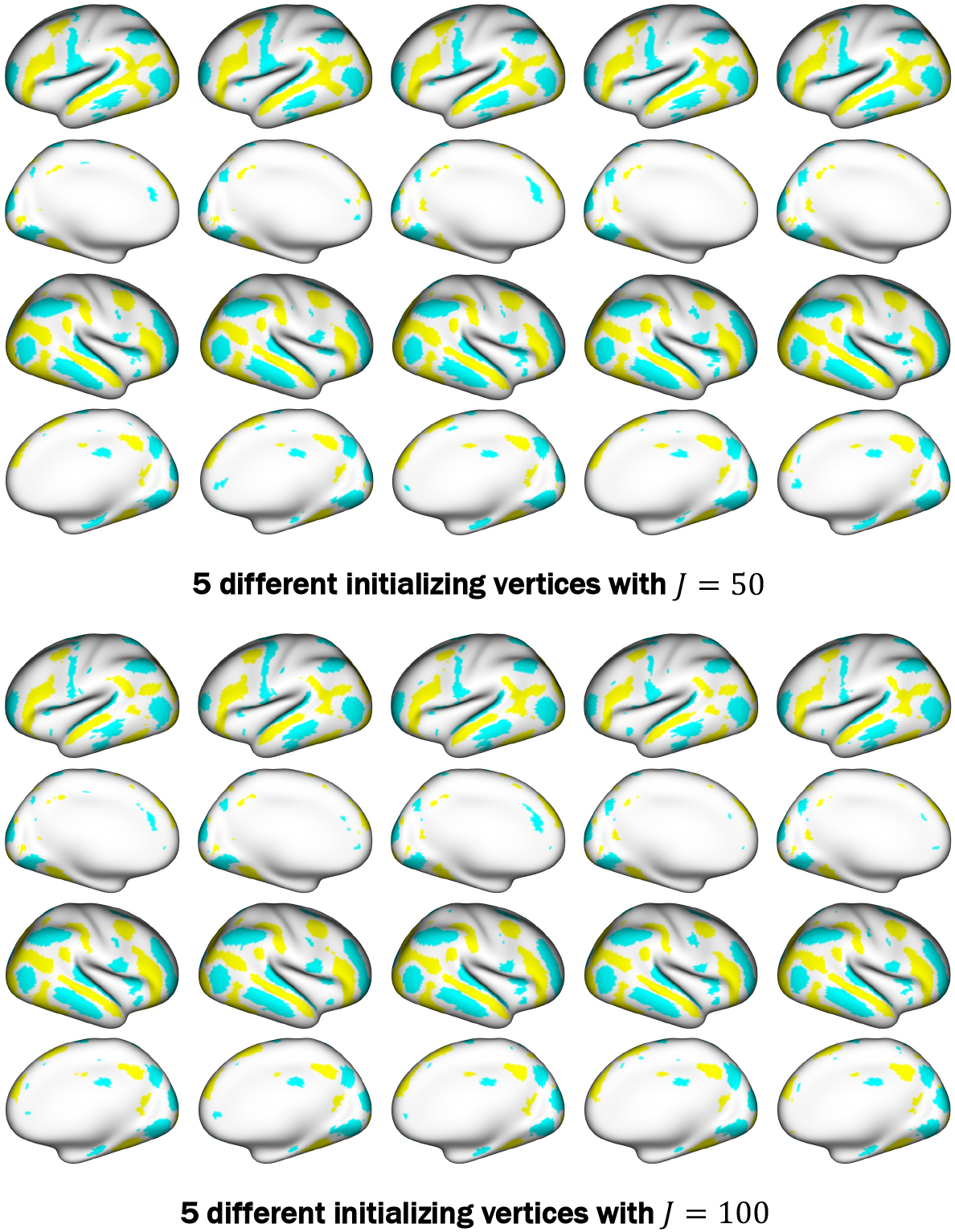
Analysis of the test data of 44 subjects with 5 different vertices that initialize the NNGP. The initializing vertex is the same for each column image.

## D. Empirical covariogram analysis

Using the contrast of parameter estimates (COPEs) data from the HCP’s social cognition and relational processing tasks, we first subtracted the group mean from each vertex, then computed the empirical covariogram for each subject. We used geodesic distance information from the spherical surface. We then computed the group-level empirical covariogram by taking the point-wise average. Figure 10 suggests that the exponential kernel closely approximates the spatial covariance of both data showing a slowly-decaying pattern.

**Figure 10:**
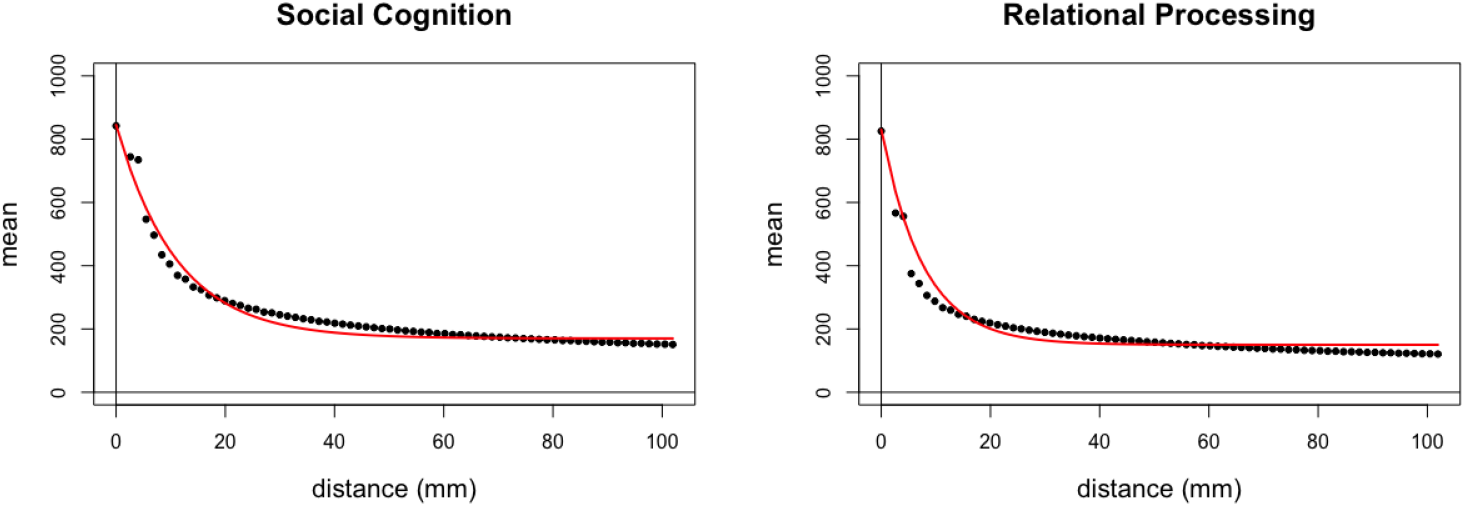
Empirical covariogram of the social cognition and relational processing tasks from the Human Connectome Project. The red line denotes for the fitted parametric covariogram using the exponential kernel.

## E. Analysis of the retest data

We used 44 subjects from the HCP’s social cognition task, who completed the task twice (test-retest). We fitted CLEAN to each of test and retest data and identified the areas of significance. As shown in Figure 11, CLEAN showed high reproducibility of the activation areas.

**Figure 11:**
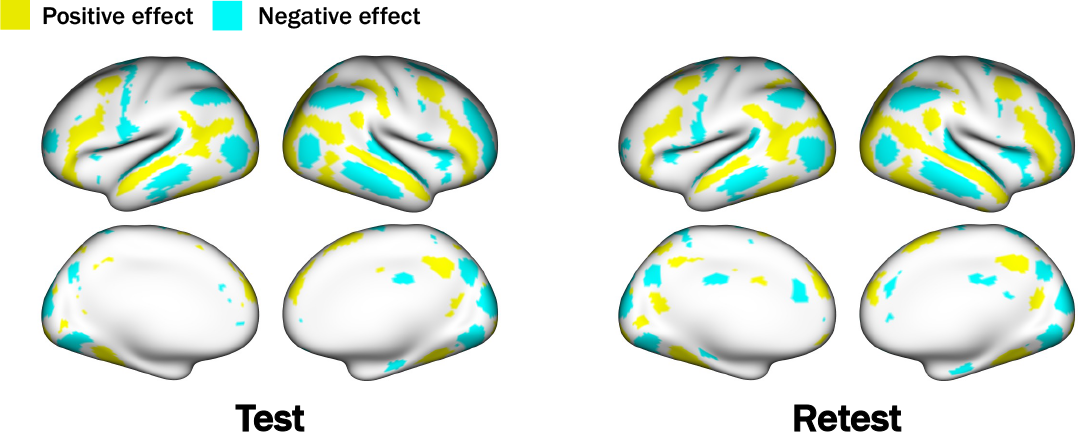
Areas of significance identified by CLEAN for both test data (left two columns) and retest data (right two columns).

## F. Simulation studies for the covariance regression

We conducted simulation studies to evaluate how the proposed method estimates parameters for spatial autocorrelation (*σ*^2^, *τ*^2^, *ϕ*). We first randomly sampled 3,000 vertices from the spherical surface from the left hemisphere. Using the exponential spatial autocorrelation function, we set two parameter sets for evaluations.

- Set 1: *σ*^2^ = 500, *τ*^2^ = 200 and *ϕ* = 0.001 (median correlation: 0.60)
- Set 2: *σ*^2^ = 200, *τ*^2^ = 500 and *ϕ* = 0.001 (median correlation: 0.25)

We generated *N* images from 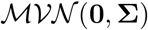, where *N* = 20, 30, 40. We then applied the proposed method to estimate these parameters (i) without any covariate (*q* = 0), (ii) with an intercept (*q* = 1), and (iii) with an intercept and a covariate generated randomly from the standard normal distribution (*q* = 2). When *q* > 0, we first regressed out the covariate effect for each vertex and used residuals to minimize the loss function. In each setting of *N* and *q*, we repeated simulations 2,000 times and took the average of the parameter estimates.

The simulation results are summarized in Table 1 and Table 2. The proposed estimator is nearly unbiased, and the standard error decreased as (i) the number of subjects increased, (ii) less covariates are used, or (ii) the proportion of the variability explained by the nugget effect *τ*^2^ increases.

**Table 1:**
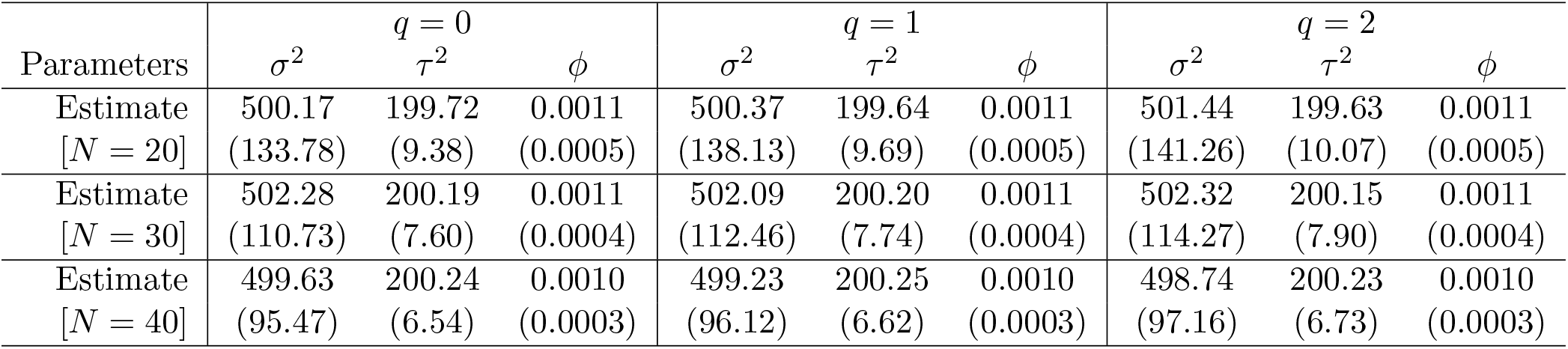
Simulation results for the parameter set 1. Parentheses denote for the standard error.

| Parameters | $q = 0$ | | | $q = 1$ | | | $q = 2$ | | |
| --- | --- | --- | --- | --- | --- | --- | --- | --- | --- |
| | $\sigma^2$ | $\tau^2$ | $\phi$ | $\sigma^2$ | $\tau^2$ | $\phi$ | $\sigma^2$ | $\tau^2$ | $\phi$ |
| Estimate<br>[$N = 20$] | 500.17<br>(133.78) | 199.72<br>(9.38) | 0.0011<br>(0.0005) | 500.37<br>(138.13) | 199.64<br>(9.69) | 0.0011<br>(0.0005) | 501.44<br>(141.26) | 199.63<br>(10.07) | 0.0011<br>(0.0005) |
| Estimate<br>[$N = 30$] | 502.28<br>(110.73) | 200.19<br>(7.60) | 0.0011<br>(0.0004) | 502.09<br>(112.46) | 200.20<br>(7.74) | 0.0011<br>(0.0004) | 502.32<br>(114.27) | 200.15<br>(7.90) | 0.0011<br>(0.0004) |
| Estimate<br>[$N = 40$] | 499.63<br>(95.47) | 200.24<br>(6.54) | 0.0010<br>(0.0003) | 499.23<br>(96.12) | 200.25<br>(6.62) | 0.0010<br>(0.0003) | 498.74<br>(97.16) | 200.23<br>(6.73) | 0.0010<br>(0.0003) |

**Table 2:** Simulation results for the parameter set 2. Parentheses denote for the standard error.

| Parameters | $q = 0$ | | | $q = 1$ | | | $q = 2$ | | |
| --- | --- | --- | --- | --- | --- | --- | --- | --- | --- |
| | $\sigma^2$ | $\tau^2$ | $\phi$ | $\sigma^2$ | $\tau^2$ | $\phi$ | $\sigma^2$ | $\tau^2$ | $\phi$ |
| Estimate<br>[$N = 20$] | 200.63<br>(53.52) | 499.85<br>(4.69) | 0.0011<br>(0.0004) | 200.06<br>(54.87) | 499.84<br>(4.78) | 0.0011<br>(0.0005) | 200.19<br>(56.44) | 499.80<br>(4.93) | 0.0011<br>(0.0005) |
| Estimate<br>[$N = 30$] | 200.91<br>(42.91) | 499.93<br>(3.78) | 0.0011<br>(0.0003) | 200.91<br>(43.75) | 499.91<br>(3.85) | 0.0011<br>(0.0004) | 201.00<br>(44.64) | 499.91<br>(3.95) | 0.0011<br>(0.0004) |
| Estimate<br>[$N = 40$] | 200.52<br>(39.81) | 500.10<br>(3.31) | 0.0010<br>(0.0003) | 200.49<br>(40.39) | 500.12<br>(3.37) | 0.0010<br>(0.0003) | 200.45<br>(40.93) | 500.12<br>(3.42) | 0.0010<br>(0.0003) |

